# The evolution of size-dependent competitive interactions promotes species coexistence

**DOI:** 10.1101/2021.05.20.445031

**Authors:** Jaime M. Anaya-Rojas, Ronald D. Bassar, Tomos Potter, Allison Blanchette, Shay Callahan, Nick Framstead, David Reznick, Joseph Travis

**Affiliations:** Department of Biological Science, Florida State University, Tallahassee Florida, USA; Department of Biology, Williams College, Williamstown Massachusetts, USA; Department of Zoology, University of Oxford, Oxford, UK; University of Illinois at Urbana–Champaign, Illinois, USA; Department of Evolution, Ecology and Organismal Biology, University of California, Riverside, CA 922521, USA

**Keywords:** size-dependent competition, intra- and interspecific competition, coexistence, coevolution, size-dependent competitive asymmetry, intraguild predation

## Abstract

1. Theory indicates that competing species coexist in a community when intraspecific competition is stronger than interspecific competition. When body size determines the outcome of competitive interactions between individuals, coexistence depends also on how resource use and the ability to compete for these resources change with body size. Testing coexistence theory in size-structured communities, therefore, requires disentangling the effects of size-dependent competitive abilities and niche shifts.
2. Here, we tested the hypothesis that the evolution of species and size-dependent competitive asymmetries increased the likelihood of coexistence between interacting species.
3. We experimentally estimated the effects of size-dependent competitive interactions on somatic growth rates of two interacting fish species, Trinidadian guppies *(Poecilia reticulata)* and killifish *(Rivulus hartii).* We controlled for the effects of size-dependent changes in the niche at two competitive settings representing the early (allopatric) and late (sympatric) evolutionary stages of a killifish-guppy community. We fitted the growth data to a model that incorporates species and size-dependent competitive asymmetries to test whether changes in the competitive interactions across stages increased the likelihood of species coexistence from allopatry to sympatry.
4. We found that guppies are competitively superior to killifish but were less so in sympatric populations. The decrease in the effects of interspecific competition on the fitness of killifish and increase in the interspecific effect on guppies’ fitness increased the likelihood that sympatric guppies and killifish will coexist. However, while the competitive asymmetries between the species changed consistently between allopatry and sympatry between drainages, the size-dependent competitive asymmetries varied between drainages.
5. These results demonstrate the importance of integrating evolution and trait-based interactions into the research of how species coexistence.

## Introduction

A persistent challenge in ecology is to understand how species can coexist when they overlap in their resource use (Chesson, 2000). Whether two such species will coexist depends upon the balance between the negative effect of competition over shared resources and the positive effect of using some resources exclusively (Chesson, 2000; Siepielski & McPeek, 2010). For many organisms, this balance depends upon the values of traits that determine the type of resource use and the efficiency with which the resources are acquired and transformed into somatic growth, survival, and offspring (Ellner *et al*., 2016).

Body size is often a key trait influencing competitive ability and resource use (Werner & Gilliam, 1984). For example, taller plants can access more sunlight and larger fish can forage more efficiently and consume a wider range of prey sizes (Weiner, 1990; Young, 2004; Bassar *et al*., 2016). In both cases, larger individuals are better competitors, a situation termed size-dependent competitive asymmetry (Weiner, 1990). In size-structured communities, size-dependent competitive asymmetries combine with competitive inequalities based on species’ identity define the differences in competitive effects between individuals on the same and other species across their respective life cycles (Bassar *et al.*, 2016).

In addition to size-based competitive asymmetries, changes in body size can be associated with changes in resource use (Hjelm *et al*., 2003; de Roos & Persson, 2013; Aresco *et al*., 2015; Turner Tomaszewicz *et al*., 2017). When this occurs, coexistence depends on how changes in body size in each species influence resource use shifts, competitive asymmetries, and fitness, particularly in the range of body sizes in which the species have the greatest overlap in resource use (Loreau & Ebenhoh, 1994; Miller & Rudolf, 2011; Nakazawa, 2015; Bassar *et al*., 2017b).

Testing theoretical predictions about coexistence in size-structured communities, however, is an empirical challenge. Compelling tests require understanding two things: first, how body size variation translates to competitive asymmetries and differences between the species in resource use; and second, how these size-based interactions map to fitness effects across the life cycles of the interacting species (Bassar *et al*., 2017b; Chesson, 2000; Siepielski & McPeek, 2010). The first requirement, separating individual effects of the confounded changes in resource use and efficiency during ontogeny, can be met with careful experimentation (Inouye, 2001; Potter *et al*., 2019).

The second requirement can only be met by integrating the demography of each species with the knowledge acquired from the first requirement (Bassar *et al*., 2017b). This is because, while competitive interactions and resource use may change with ontogeny, the various stages in the life cycle do not count equally towards fitness (Caswell, 2001). Reproductive value (percapita contribution of age- or size-class towards fitness), for instance, can change dramatically as organisms grow or age (Caswell, 2001; Bassar, Travis & Coulson, 2017b)

A solution to these challenges is to develop demographic models of interacting structured populations that can be readily parameterised by empirical data (Ellner *et al*., 2016). These models allow researchers to evaluate how changes in the ecological interactions between the species influences their fitness (Bassar *et al*., 2016, 2017b,a). Applying such models requires devising manipulative experiments that can measure, separately for each species, how trait variation influences resource use and competitive efficiency, and ultimately, the fitness consequences of trait variation integrated across the life cycles of the organisms.

Here, we experimentally test how species- and size-dependent competitive asymmetries contribute to the evolution of species coexistence in Trinidadian stream communities comprised of Trinidadian guppy *(Poecilia reticulata)* and Hart’s killifish *(Rivulus hartii)* a.k.a., KG communities (Travis *et al.*, 2014). We executed a laboratory experiment that allowed us to parameterise a size-structured demographic model for these species under two contrasting scenarios, representing the beginning and endpoint of novel community formation (Travis *et al*., 2014; Bassar *et al*., 2017a). Prior empirical work has demonstrated that coexistence between these species is an evolved property, with coexistence more likely between established sympatric populations of each species than between allopatric populations that first encounter each other (Bassar *et al*., 2017a). However, it is not clear whether the increased likelihood of coexistence is due to changes in competitive asymmetries, changes in the resource niche use, or both. Specifically, we tested the hypothesis that the evolution of stronger size-based competitive asymmetries in guppies increased the likelihood of coexistence between both species in sympatry.

## Methods

### Study system

Each river that drains the Northern Mountain Range of the Caribbean island of Trinidad has a replicated succession of fish communities. At low elevations, fish communities contain numerous species (Gilliam *et al*., 1993). Fish species diversity declines progressively upstream because waterfalls impede the upstream movement of larger fish (Gilliam *et al*., 1993). In lower stream reaches, guppies and killifish co-occur with multiple predatory fish species (killifish-guppy-predator communities, hereafter KGP communities). Above barrier waterfalls, guppies and killifish occur without predators (killifish-guppy, or KG communities). Above these communities, killifish are the only fish species found in the streams (killifish only, or KO communities). Life histories, behaviour, morphology, and physiology evolve in both species as they adapt to these different communities (Reznick & Endler, 1982; Ghalambor et al., 2004; Auer *et al*., 2018).

KG communities are thought to be formed when guppies are able to surmount barrier waterfalls and invade KO communities (Travis *et al.*, 2014). This encounter between KGP guppies and KO killifish represents the first stage of KG community development with allopatric phenotypes. Replicated experiments, wherein guppies from KGP communities were translocated over barrier waterfalls into KO communities have shown that both species evolve genetically based trait differences that are consistent with those observed in comparative analyses of natural KG communities (Reznick *et al.*, 1990, 2019; Walsh & Reznick, 2011). These evolutionary changes have been observed over relatively short time periods (Reznick *et al*., 2019). In this study, we consider sympatric phenotypes of killifish and guppies to be those from long-established KG communities that have evolved together (Alexander *et al.*, 2006; Walter et al., 2011). These phenotypes represent the endpoint of the evolutionary interaction between the species in these locations (Bassar et al., 2017a).

Guppies and killifish use similar resources, which makes their coexistence a persistent puzzle. There is some evidence suggesting that both fish species alter their resource use after guppies have invaded KO streams. Guppies feed primarily on aquatic invertebrates, detritus, and algae (Bassar *et al.*, 2010; Zandona *et al.*, 2011; Fraser & Lamphere, 2013) but guppies from KGP communities feed mostly on invertebrates, while guppies in KG communities are more generalist feeders (Zandona *et al*., 2011). Moreover, dietary studies based on stomach contents have shown little support for strong size-based niche shifts in guppies (Zandonà *et al*., 2015). Killifish, on the other hand, are mostly insectivorous (Fraser & Gilliam, 1992) and are thought to have stronger size-based niche shifts than guppies (Travis *et al*., 2014). Field observations have shown that killifish prey on aquatic and terrestrial invertebrates (Owens, 2010; Murray et al., 2018). But there is no evidence of strong size-based niche shifts in killifish.

Theory shows that different patterns of size-based niche use between the species can lead to coexistence only if guppies are better competitors than killifish in the region of the niche that is shared between the species (i.e., aquatic invertebrates; Bassar, Travis & Coulson, 2017b). Mark-recapture and experimental studies have shown that larger guppies are better intraspecific competitors than smaller guppies and that guppies from KG communities are generally stronger intraspecific competitors than KGP guppies (Bassar *et al.*, 2013; Potter et al., 2019).

### Demographic model

Our experiment was designed to estimate the parameters describing the interactions between individuals of distinct species and body sizes, parameters that feed directly into a size-structured demographic model (Bassar, Travis & Coulson, 2017b). A model that allows us to map competitive interactions across the life cycle onto the fitness of both species. The demographic model is an integral projection model (IPMs) described in Bassar et al. (2016) and Bassar, Travis & Coulson (2017b). A key advantage of these type of models is that they can be parameterised with individual data on demographic rates, allowing the generation of theoretical predictions and interpretation of experimental data (Bassar *et al*., 2016, 2017b; Potter *et al*., 2019). We briefly summarise the salient features of the model here. More details can be found in Bassar et al. (2016) and Bassar, Travis & Coulson (2017b).

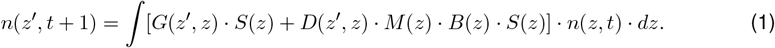

where *n*(*z,t*) is a distribution describing the number of individuals of body size *z*’ at time *t, n*(*z’,t* + 1) is a distribution describing the number of individuals of *Z* at time *t* + 1. The functions *S, G, B, M,* and *D* are demographic rates representing survival, growth, probability of reproduction, litter size, and size of offspring in individuals as functions of body size *z* at time *t*.

If *V_i_* = [*S_i_, G_i_, B_i_, M_i_, D_i_*] is the set of the linearised demographic rate equations for species *i, z_i_* is the body size of a focal individual of species *i*, and *x_i_* represents the body sizes of intraspecific competitors, then an equation for any of the mean demographic rates of species *i* competing with species *j* and other members of species *i* can be expressed as:

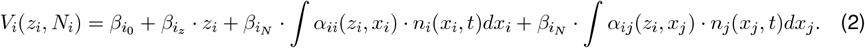

The first two terms (*β*_*i*_0__ + *β_i_z__* · *z_i_*) describe the value of the demographic rate of an individual of species *i* and size *z* in the absence of competition. The parameter *β_i_N__* describes the per-capita effect on the demographic rate *V* (e.g., somatic grow rates or survival) of an individual of species *i* and size *z* of an equal sized individual of the same species. The function *n*(*x, t*) describes the distribution of sizes of competitors at time *t* and may be either from the same species, *i*, or from another species, *j*. The species- and size-dependent effects of intraspecific and interspecific competition on the demographic rates are calculated as interaction surfaces *α_ii_*(*z_i_,x_i_*) and *α_ij_*(*z_i_,x_j_*), respectively, which describe the competitive equivalence of individuals of size *x_i_* or *x_j_* on individuals of size *z_i_*. Overall, the interaction surfaces can be interpreted as the relative competitive effect of a competitor of species *i* (or *j*) of size *x* on an individual of size *z* of species *i*. For example, if *α_ij_*(*z_i_,x_j_*) = 3, it means that the competitive effect of a competitor of size *x* of species *j* on a size *z* individual of species *i* is equivalent to the competitive effects of three individuals of species *i* and size *z* (see Bassar, Travis & Coulson, 2017b).

The outcome of competition can be determined from the shape of the interaction surfaces. In turn, the shape of the interaction surface depends upon the resources that are used by each species and on the size-dependent ability of individuals to effectively compete for those resources. When individuals use different resources and larger or smaller individuals are better at acquiring and assimilating those resources, the interspecific interaction surfaces can be modelled as an exponential function of the size and species of competitors as:

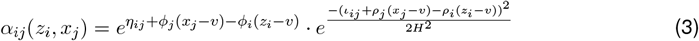

The expression 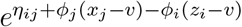 describes how the competitive effects change as a function of species identity and body size (i.e., the species-dependent and size-dependent competitive asymmetry component). The expression 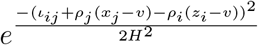 describes the proportion of niche overlap between individuals of each species as a function of their body sizes.

In the first expression, the parameter *η_ij_* captures the relative species-dependent competitive effect, which is simply the difference between the competitive effects of intraspecific competitors (species *i)* and interspecific competitors (species *j*) at size *v*, which is a centring value chosen by the researcher. Typically, *v* is a biologically meaningful size, such as size at birth. If *η_ij_* < 0, the competitive effect of species *j* on species *i* at size *v* is less than the competitive effect of species *i* is on itself. The size-dependent competition coefficients *ϕ_i_* and *ϕ_j_* describe how the magnitude of the effect changes with body size in each species (i.e., size-dependent competitive asymmetry). When *ϕ_i_* = 0, the competitive effects of species *i* are said to be symmetric with respect to body size of species *i* (i.e., body size plays no role in determining the competitive advantage of species i). When *ϕ_i_* > 0, competition is considered asymmetric with respect to size, and larger individuals have stronger competitive effects than smaller individuals. Conversely, when *ϕ*_i_ < 0, competition is asymmetric with respect to size such that smaller individuals have stronger competitive effects than larger individuals. Overall, the *η* and *ϕ* parameters together describe the relative competitive effects of one species on the other. The degree of interspecific competitive asymmetry is therefore given by the difference between the effects of conspecific competitors on focal growth (across all sizes) and the effects of heterospecific competitors on focal growth (across all sizes).

Following previous treatments of niche overlap (MacArthur & Levins, 1967), the resource niche is modelled as a normal distribution along a resource axis, *R* (Bassar, Travis & Coulson, 2017b). The niche overlap, as a function of body size, is the overlap of the distributions of individuals with trait value *z* from species *i* and trait value *x* from species *j*. The parameter *ι_ij_* is the difference in the mean of the resource niche at size *v*; and *ρ_i_* and *ρ_j_* describe how the mean of the resource niche changes with body size in species *i* and *j*, respectively. *H*^2^ is the variance in the niche width. In our experiment described below, we assume that *ι_j_*, *ρ_i_*, and *ρ_j_* are zero. In this case, the niche overlap term is unity for all sizes, meaning all individuals compete for the same resource. When the experiment is carried out using a single resource – as in this study– this allows researchers to fit the demographic model to the empirical data without confounding changes in competitive effects with changes in resource use.

### Experimental design

We were interested in asking whether there is evidence for differences in species- and size-dependent competitive effects (*η* and *ϕ*’s) for killifish and guppies between the initial (allopatric) and final stages (sympatric) of community development and whether these changes would lead to a greater likelihood of coexistence. We estimated these parameters by performing an aquaria-based response surface experiment (Inouye, 2001), in which we manipulated the number and size distributions of fish in the treatments (Fig. 1). We measured somatic growth over a 28-day period and fitted a version of equation 2 to those data. Because theory predicts that the two species could coexist if guppies are better competitors for resources on the shared portion of the niche (i.e., aquatic invertebrates), we fed the fish identical quantified rations of aquatic invertebrates over the course of the experiment. We ran this experiment using guppies and killifish with allopatric phenotypes (KO killifish and KGP guppies) and sympatric phenotypes (KG guppies and KG killifish). The pairing of KO killifish with KGP guppies recreates the initial stages of the invasion of a KO community by guppies from KGP communities. The pairing of KG killifish with KG guppies represents the situation after each species has adapted to the other.

**Figure 1.**
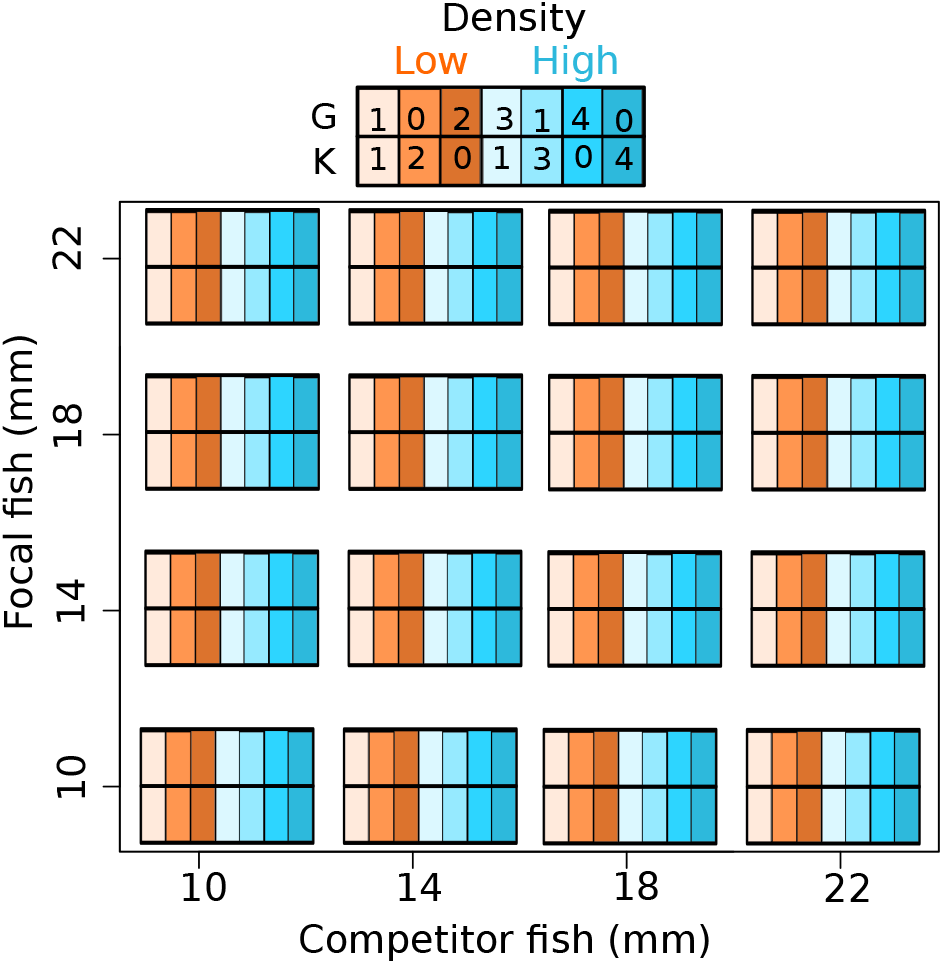
Schematic representation of the design of the experiment. Each coloured cell represents an aquarium, in which we manipulated the density (high and low: orange and blue cells, respectively) and frequency of guppies (G) and killifish (K) across a range of body sizes. Within each aquarium, each fish acted as a competitor (X-axis) and as a focal fish (y-axis).

Each aquarium contained either two or four fish (Fig. 1). Within the two fish treatment, the tanks contained either two guppies, two killifish, or one guppy and one killifish. Within the four fish treatment, the tanks contained either four guppies, four killifish, three guppies and one killifish, or one guppy and three killifish. Manipulating the sizes of fish in each tank produced different combinations of competitor sizes and enabled the experiment to cover different portions of the interaction surfaces. All fish were classified into four size classes (±2*mm*): 10*mm*, 14*mm*, 18*mm*, and 22*mm*. These size classes represent different life-history stages in guppies (10 and 14*mm* = juveniles, 18*mm* = young adults, 22*mm* = older adult females; Reznick *et al*., 2001) and juvenile stages of killifish, which mature at approximately 35*mm* (Walsh & Reznick, 2008). Each aquarium had either one or two size classes of fish, and sizes were paired so that each size category competed against fish of all other size categories in the experiment (112 possible unique competition trials). Simulations and experimental studies have shown that this experimental design has enough power to estimate the parameters of the interaction surfaces (Bassar *et al*., 2017b; Potter *et al*., 2019). Some experimental trials were not possible because 22*mm* killifish tended to kill guppies smaller than 18*mm*.

### Sampling and allocating fish to treatments

We ran the experiment with fish from two different river systems, the Aripo and the Quare. These systems represent independent origins of allopatric KGP guppies invading KO habitats to establish coevolved sympatric KG communities (Willing *et al*., 2010; Walter *et al*., 2011). In the Aripo drainage, we collected guppies from KGP communities downstream from Haskins’ Falls on the Aripo River. We collected KG guppies and killifish from the Naranjo and Endler streams and KO killifish from the upper reaches of the Naranjo stream. In the Quare drainage, we collected KGP guppies from the main branch of the Quare River, adjacent to the pump-house on the Hollis Reservoir Road. We collected KG guppies and killifish from the El Campo and Quare 2 tributaries to the Quare River and KO killifish from the upper reaches of each of these tributaries.

We performed our experiments during the dry season (February to June). We collected guppies with butterfly nets and transported them to our field station in plastic bottles with medicated water (0.15*mL/L* of Stress Coat^®^, Mars Fishcare, PA, USA; 0.075*mL/L* AmQuel Plus^®^, Kordon LLC, CA, USA). Killifish were collected with hand nets and transported to the laboratory in Hefty^®^-bags. In the laboratory, we treated all fish with a salt bath (sea salt 25*g/l*, 15*min*) to eliminate ectoparasites and with antibiotics (0.187*g*/20*L* Tetracycline or 1.25*g*/20*L* Furan) to reduce the likelihood of bacterial infections.

At the beginning of the experiments, we assigned each fish to an appropriate size class based on standard length (SL, ±0.5*mm*), measured its mass (±0.001*g*), and classified it as male, female, or juvenile. We used only females and juvenile guppies, and juvenile killifish in our experiment. We did not use adult male guppies because they have little or no growth after maturity (14*mm*), which limits our ability to evaluate the fitness consequences of the competitive environment. We randomly assigned each individual to a size class and density treatment (Fig. 1). Before adding them to their tanks, we marked individuals with a single subcutaneous injection of a coloured elastomer (Northwest Marine Technology) for identification. We kept extra fish in glass tanks at approximately two fish per litre of water to replace any fish that died during the experiment, and to maintain the density and size-structure treatment for the experimental fish.

### Fish feeding and housing

We performed the competition trials in plastic tanks (*L* · *W* · *H*: 26 · 16 · 17*cm*) for 28 days and provided food to each tank once a day as a 1000 *μl* solution of *Artemia* nauplii (≈ 400 live larvae). We filled each tank with ≈ 4*L* of medicated water (0.15*mL/L* of Stress Coat, Mars Fishcare, PA, USA; 0.075*mL/L* AmQuel Plus^®^, Kordon LLC, CA, USA). On the first day of the experiment, we added 500*μL* of 10% Formalin to each tank to further reduce ectoparasites such as *Gyrodactylus spp.* and to avoid secondary bacterial infections. We performed water changes every other day by siphoning out up to 75% of the water, making sure to remove organic waste and excess food. Dead or missing individuals were replaced with another fish of the same species, community of origin, and size class to maintain the experimental treatment.

### Data analysis

#### Statistical model

We estimated the competition parameters in equation 2 using the somatic growth increment as a demographic rate (*V*). The somatic growth increment is an ideal demographic rate from which to estimate the effects of competition –both fish species of these size classes grow considerably over a 28-day period – and fitness is extremely sensitive to effects on somatic growth (Bassar *et al*., 2013, 2017a). The demographic equation for somatic growth is a slightly modified version of equation 2 (Griffiths *et al*., 2020). Our somatic growth increment equation for guppies was:

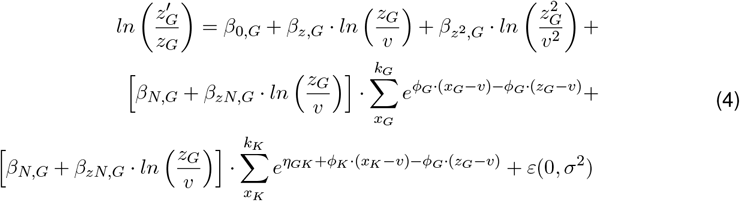

Where 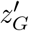 is the observed length of an individual guppy after the interval and *z_G_* is the observed length of the same guppy at the beginning of the experiment. The somatic growth increment typically decreases with increased length (*β*_*zG*_) and sometimes is slightly convex, which is captured by the quadratic term, *β*_*z*^2^,*G*_. We also included other terms to the model (*β_N,G_* and *β_zN,G_*) that describes how individuals of distinct sizes respond to resource competition. The integrals in equation 2 are replaced by summations in equation 4 because the populations in the experiment are small and are better described in discrete terms rather than continuous functions. The indexes on the summation thus represent individual guppies or killifish. The term *ε*(0, *σ*^2^) is a normal residual error term with mean of zero and variance *σ*^2^. An analogous equation was fitted to the killifish data.

#### Model fitting

We estimated all parameters in equation 4 using a Bayesian modelling framework in Stan via the *rstan* package in R 3.6 (Carpenter *et al*., 2017). We fitted data for the Quare and Aripo drainages separately. To allow comparisons between species and phenotypes within drainages, we fit models that included both species from allopatric and sympatric locations. We used dummy coding to identify parameters for the distinct species and phenotypes. Posteriors were sampled from six Hamilton Monte Carlo (HMC) chains, 8000 iterations, and a warmup of 5000 iterations. We used informative priors for the guppy interaction surface parameters that we derived from similar experiments on intraspecific competition using KGP and KG guppies (Potter *et al*., 2019). For the rest of the parameters, we used weakly informative priors, e.g., mean= 0 and SD= 5. For all models, we verified that all four chains converged using the estimated potential scale reduction statistic 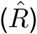 and visually checking the trace plots (Fig. S1-S6). Additionally, we evaluated each model’s behaviour by plotting the predicted vs observed values and the distribution of their residuals (Fig. S7).

We modelled somatic growth increment with a normal error distribution and length centred at 10*mm* (i.e., *v*). To facilitate the interpretation of the parameters describing the interaction surface, beginning from near birth to older individuals, we set *v* equal to 10*mm* in both species. This estimates the difference between the species in their competitive abilities at 10*mm*. We report the means of the posterior samples of the parameters and the 95% Compatibility Interval (i.e., mean [2.5%, 97.5% CI]). For comparisons between parameters, we estimated the difference between the posterior samples and reported the mean differences, CI, and the Level of Support (LOS) of the difference in the parameters. The LOS is estimated as the proportion of the posterior difference distribution greater or less than zero. We asked whether the species and size-dependent competition parameters improved the overall fit of the growth models to the data by comparing the models for each treatment with a null model assuming symmetric species and size-dependent competition parameters (*η_GK_* = 0; *ϕ_G_* = *ϕ_K_* = 0) using the function *compare()* from the rethinking R package (McElreath, 2020). We chose the best model as the model with the lowest WAIC (Watanabe Akaike Information Criterion) and highest weight.

For the statistical analyses, we used the growth data of any fish that began and completed the experiment (776/891; 87%). Survival to the end of the experiment was higher in the Aripo compared to the Quare (92% vs 82%), yielding slightly higher sample sizes in the Aripo compared to the Quare. We replaced any fish that died during the experiment with a similar-sized fish. These replacements were not used as dependent data for growth unless they were in the experiment for more than 25 of the 28 days of the experiment. We used the size of the replacement fish to calculate the weighted average size of competitors. To do so, we multiplied the body size of the replaced and replacement fish by the number of days they were in the experiment and divided by the total number of days in the experimental tanks.

#### Comparisons of the change in the competitive asymmetries between communities

We used the posterior samples of the parameters describing the interaction surfaces to ask how the differences between the intraspecific and interspecific competitive effects of each species on the other changed between allopatric (i.e., KGP guppies vs KO killifish) and sympatric phenotypes (i.e., KG guppies vs KG killifish). For each species, we calculated the change in competitive asymmetry by subtracting the difference in the intra- and interspecific interaction surfaces from the allopatric trials (representing the initial conditions following the invasion of guppies) from the difference in the intra- and interspecific interaction surfaces in the sympatric trials (representing the competitive conditions in established KG communities). For example, for guppies, we calculated the change in competitive asymmetry as (Δ*CA*):

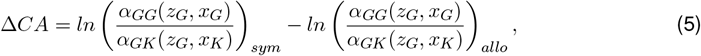

where subscripts “*sym*” and “*allo*” denote the sympatric and allopatric comparisons, respectively. The effect is independent of the size of the individual experiencing the competitive impact (i.e., *z_G_*), which can be seen by replacing the alphas with the parameters 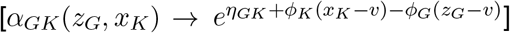 and rearranging them:

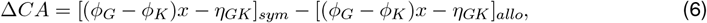

Calculating the difference in competitive asymmetries in this way captures how the differences between the effects of intraspecific and interspecific competitors change from allopatric to sympatric phenotypes. Positive values of Δ*CA* indicate that the effects of interspecific competition are stronger in sympatric phenotypes compared with allopatric ones, negative values of Δ*CA* indicate that the effects of interspecific competition are weaker in sympatric phenotype, compared with allopatric ones.

#### How changes in competitive asymmetry alter predictions of coexistence

We parameterised the model of Bassar, Travis & Coulson (2017b) using the posterior means of the parameter from this study to illustrate how changes in the interaction surfaces between the two communities alter predictions about species coexistence. We parameterised the model, assuming that the only differences between the species are the values of the intra- and interspecific interaction surfaces. All other parameters were identical and based on KG guppies, as described in Bassar, Travis & Coulson (2017b). Using parameters from a single species and changing only the competition parameters isolates the effect of the change in the competition parameters on coexistence. Other differences between the species that may contribute to coexistence or competitive exclusion will not be included.

We used the model to calculate the invasion exponent of each species in each community type as the dominant eigenvalue of the matrix approximation of the continuous model (for details on these calculations, Bassar, Travis & Coulson, 2017b). Coexistence is predicted by mutual invasibility, meaning that each species can invade a population of the other when the resident population is at its single species equilibrium (Siepielski & McPeek, 2010; Bassar *et al*., 2017b). We illustrate the predictions of the model by evaluating mutual invasibility over a range of ontogenetic niche shift parameters (*ρ*) in both species. We varied *ρ* from no niche shift (*ρ* = 0) to a value indicating moderate niche shifts (*ρ* = 0.18) between new-born individuals (≈ 6*mm* SL) and larger individuals (≈ 25*mm* SL), as in Bassar, Travis & Coulson (2017b).

## Results

### Are competitive interactions between guppies and killifish asymmetric and size-structured?

Yes, resource competition within and between guppies and killifish depends strongly on the species identity of their competitors (*η*) and on body size (*ϕ*, Table 1). Incorporating the parameters that describe species and size-dependent competition into the growth models increased the fit of the models to the data compared with models that did not include these effects (Δ*WAIC* > 10.9 for all four models, Table 2). At 10*mm* (SL), there was strong and consistent support for killifish exerting weaker species-dependent competitive effects on guppies than guppies on themselves (all *LOS*_*ηGK*<0_ > 97%, see Table 1 and Fig. 2b). At the same time, small (10*mm* SL) guppies exerted a stronger competitive effect on killifish than a killifish of equal size on themselves (-*η_GK_* = *η_KG_* > 0). For both species, larger individuals were stronger competitors (Table 1, Fig. 2a: *ϕ* > 0, i.e., positive size-dependent competitive effects for both species). The level of support for positive size-dependent competitive asymmetry (*ϕ* > 0) was larger than 94% in all but allopatric killifish from the Aripo drainage (*LOS*_*ϕK*>0_ = 88.4%, Table 1).

**Figure 2.**
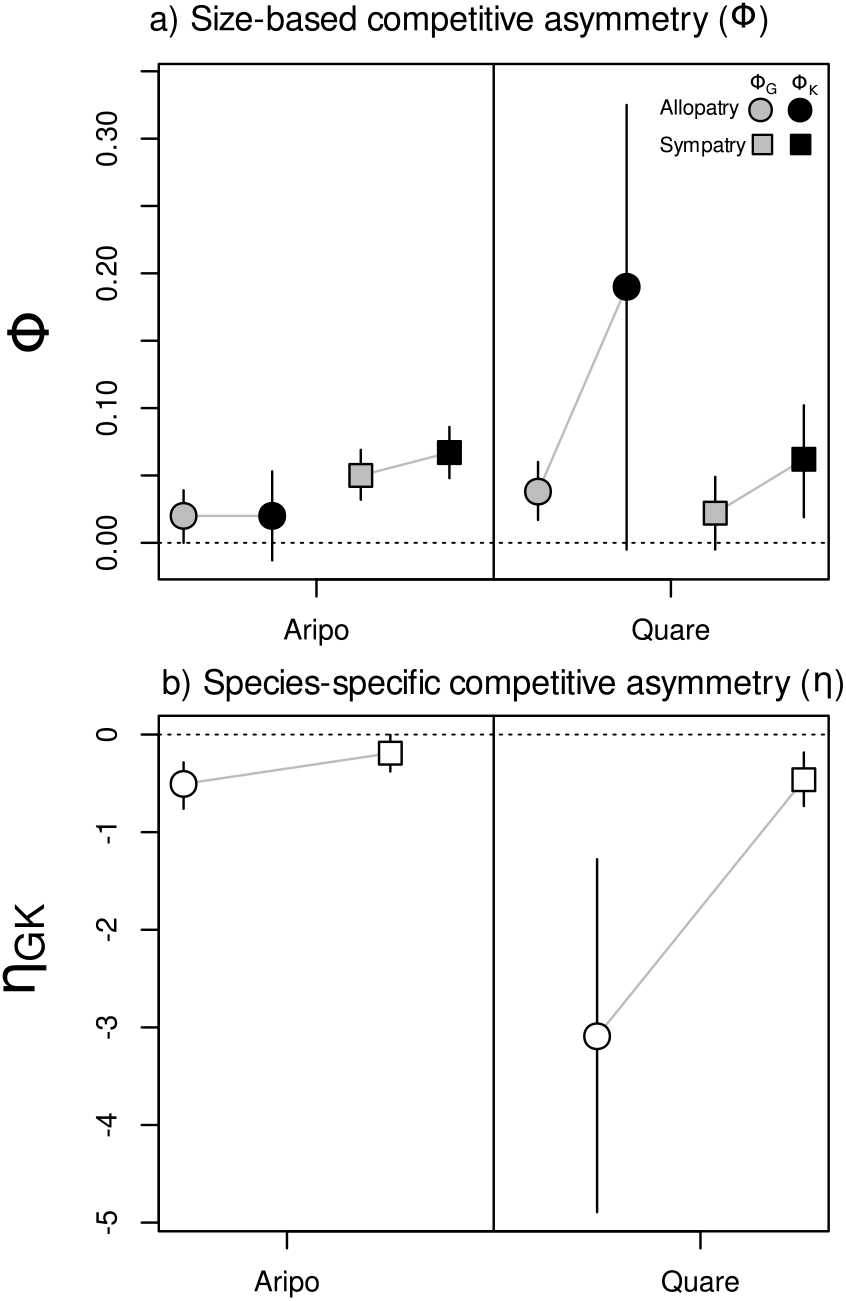
Estimated species- and size-dependent competitive asymmetry parameters. In a), values greater than zero indicate positive size-based competitive asymmetry. In b), *η_GK_* indicate that individual killifish have a smaller competitive effect on guppies than guppy of them selfs at 10*mm*.

**Table 1:** Parameter estimates of the models.

| Parameter | Aripo |  |  |  |  |  |  |  | Quare |  |  |  |  |  |  |  |
| --- | --- | --- | --- | --- | --- | --- | --- | --- | --- | --- | --- | --- | --- | --- | --- | --- |
|  | Allopatric |  |  |  | Sympatric |  |  |  | Allopatric |  |  |  | Sympatric |  |  |  |
|  | mean | SD | 2.50% (CI) | 97.50% (CI) | mean | SD | 2.50% (CI) | 97.50% (CI) | mean | SD | 2.50% (CI) | 97.50% (CI) | mean | SD | 2.50% (CI) | 97.50% (CI) |
| **Guppy** |  |  |  |  |  |  |  |  |  |  |  |  |  |  |  |  |
| $\beta_0$ | **0.397** | 0.03 | 0.337 | 0.456 | **0.328** | 0.037 | 0.255 | 0.401 | **0.423** | 0.024 | 0.377 | 0.47 | **0.318** | 0.034 | 0.25 | 0.385 |
| $\beta_z$ | **-0.762** | 0.073 | -0.909 | -0.619 | **-0.575** | 0.096 | -0.761 | -0.387 | **-0.859** | 0.068 | -0.994 | -0.727 | **-0.591** | 0.087 | -0.761 | -0.413 |
| $\beta_{z^2}$ | **0.355** | 0.054 | 0.25 | 0.462 | **0.253** | 0.068 | 0.12 | 0.386 | **0.385** | 0.058 | 0.271 | 0.499 | **0.279** | 0.087 | 0.107 | 0.45 |
| $\beta_N$ | **-0.073** | 0.009 | -0.089 | -0.056 | **-0.038** | 0.01 | -0.057 | -0.019 | **-0.081** | 0.007 | -0.095 | -0.068 | **-0.034** | 0.011 | -0.056 | -0.012 |
| $\beta_{zN}$ | **0.084** | 0.016 | 0.054 | 0.115 | **0.035** | 0.02 | -0.002 | 0.074 | **0.107** | 0.013 | 0.081 | 0.132 | **0.028** | 0.022 | -0.017 | 0.071 |
| $\sigma$ | **0.04** | 0.003 | 0.035 | 0.046 | **0.055** | 0.004 | 0.048 | 0.063 | **0.037** | 0.003 | 0.032 | 0.042 | **0.046** | 0.004 | 0.039 | 0.055 |
| **Killifish** |  |  |  |  |  |  |  |  |  |  |  |  |  |  |  |  |
| $\beta_0$ | **0.39** | 0.033 | 0.324 | 0.455 | **0.695** | 0.032 | 0.631 | 0.758 | **0.17** | 0.028 | 0.119 | 0.232 | **0.559** | 0.055 | 0.451 | 0.667 |
| $\beta_z$ | **-0.707** | 0.076 | -0.861 | -0.559 | **-0.976** | 0.09 | -1.153 | -0.799 | **-0.322** | 0.123 | -0.573 | -0.091 | **-0.863** | 0.148 | -1.153 | -0.569 |
| $\beta_{z^2}$ | **0.249** | 0.068 | 0.114 | 0.382 | **0.418** | 0.069 | 0.283 | 0.552 | **0.157** | 0.125 | -0.081 | 0.402 | **0.295** | 0.095 | 0.106 | 0.482 |
| $\beta_N$ | **-0.06** | 0.009 | -0.078 | -0.042 | **-0.096** | 0.009 | -0.114 | -0.079 | -0.002 | 0.003 | -0.009 | 0.000 | **-0.085** | 0.014 | -0.113 | -0.058 |
| $\beta_{zN}$ | **0.078** | 0.016 | 0.047 | 0.109 | **0.06** | 0.017 | 0.027 | 0.093 | -0.007 | 0.006 | -0.019 | 0.006 | **0.085** | 0.029 | 0.029 | 0.144 |
| $\sigma$ | **0.038** | 0.003 | 0.032 | 0.045 | **0.048** | 0.004 | 0.042 | 0.056 | **0.062** | 0.005 | 0.054 | 0.073 | **0.053** | 0.005 | 0.044 | 0.065 |
| **Interaction surface** |  |  |  |  |  |  |  |  |  |  |  |  |  |  |  |  |
| $\phi_G$ | 0.02 | 0.01 | 0.000 | 0.039 | **0.05** | 0.009 | 0.032 | 0.069 | **0.038** | 0.011 | 0.017 | 0.06 | 0.022 | 0.014 | -0.005 | 0.049 |
| $\phi_K$ | 0.02 | 0.017 | -0.013 | 0.053 | **0.067** | 0.01 | 0.048 | 0.086 | 0.19 | 0.082 | -0.005 | 0.325 | **0.062** | 0.021 | 0.019 | 0.102 |
| $\eta_{GK}$ | **-0.506** | 0.121 | -0.761 | -0.285 | **-0.191** | 0.096 | -0.379 | -0.003 | **-3.091** | 0.927 | -4.893 | -1.277 | **-0.463** | 0.139 | -0.732 | -0.184 |

**Table 2:**
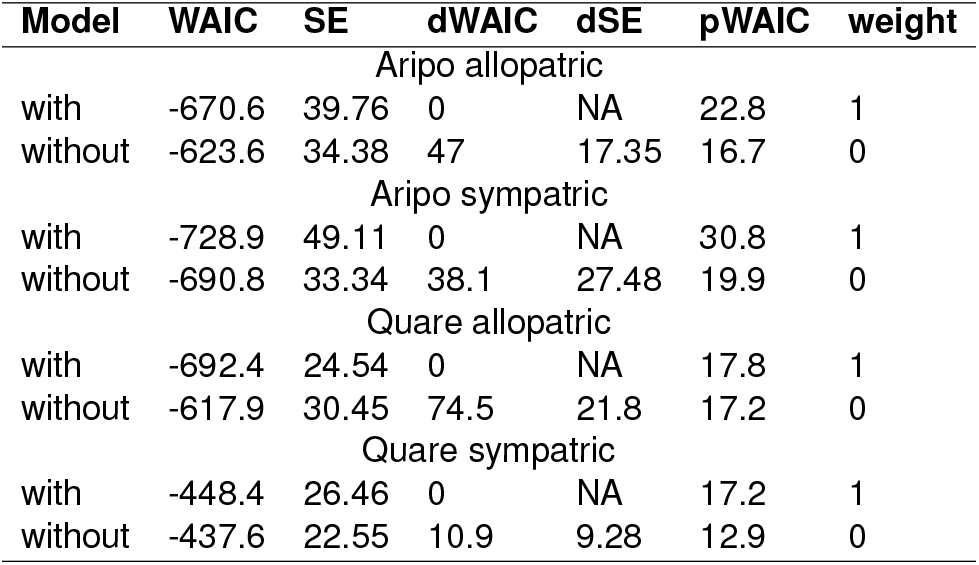
Fit statistics for growth models of guppies and killifish. Including the interaction surface parameters increased model fit (lower WAIC) and model weight.

| **Model** | **WAIC** | **SE** | **dWAIC** | **dSE** | **pWAIC** | **weight** |
| --- | --- | --- | --- | --- | --- | --- |
| Aripo allopatric |  |  |  |  |  |  |
| with | -670.6 | 39.76 | 0 | NA | 22.8 | 1 |
| without | -623.6 | 34.38 | 47 | 17.35 | 16.7 | 0 |
| Aripo sympatric |  |  |  |  |  |  |
| with | -728.9 | 49.11 | 0 | NA | 30.8 | 1 |
| without | -690.8 | 33.34 | 38.1 | 27.48 | 19.9 | 0 |
| Quare allopatric |  |  |  |  |  |  |
| with | -692.4 | 24.54 | 0 | NA | 17.8 | 1 |
| without | -617.9 | 30.45 | 74.5 | 21.8 | 17.2 | 0 |
| Quare sympatric |  |  |  |  |  |  |
| with | -448.4 | 26.46 | 0 | NA | 17.2 | 1 |
| without | -437.6 | 22.55 | 10.9 | 9.28 | 12.9 | 0 |

### Is there evidence for the evolution of changes in the competitive asymmetry between guppies and killifish following guppy invasion?

Yes, for both guppies and killifish, the level of interspecific competitive asymmetry was smaller between sympatric phenotypes – representing coadapted communities-than between allopatric phenotypes –representing the initial invasion of guppies into killifish populations (*LOS*_Δ*CA*<0_ > 99.3%, Fig. 3a and 4a). Contrary to the Aripo system, in the Quare system the level of interspecific competition increases with the size of competitors, suggesting that sympatric killifish outside the body size range of guppies have stronger competitive effects than guppies. Although the greatest change occurred in the largest size classes of competitors (middle and right columns in Fig. 3 and 4), the change in the interspecific differences in competitive asymmetry was consistent across all sizes.

**Figure 3.**
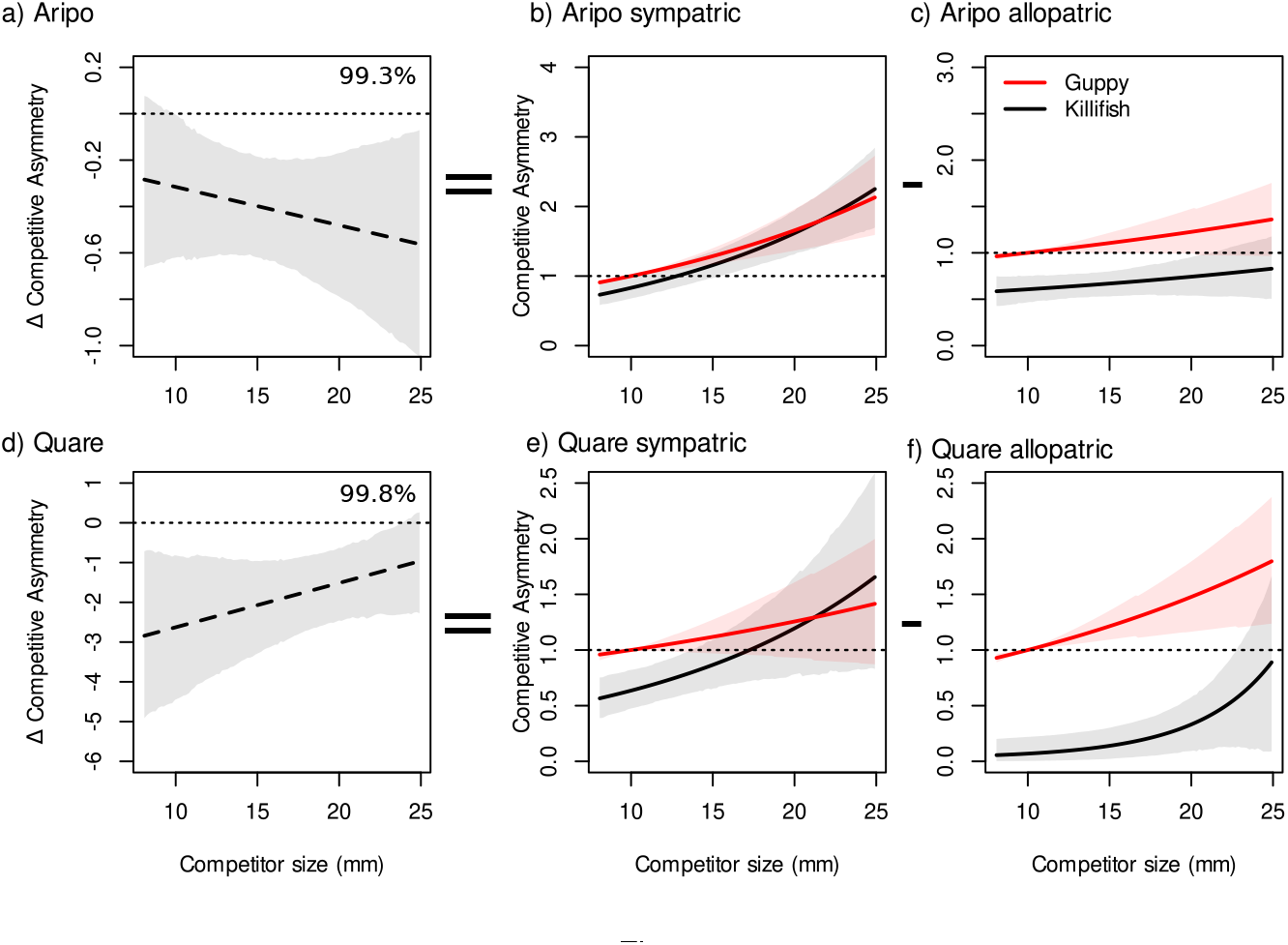
Change in competitive asymmetry for guppies. Panel a) shows the change in competitive asymmetry from allopatry to sympatry. Panels b), c), e), and f) show slices of the interaction surface for a “focal” 10mm guppy competing against either other guppies (red) or killifish (black) of all sizes. Values less than 1 mean that the competitive asymmetry favours the 10*mm* guppy, and values greater than one means favours the competitor. Shaded areas represent the 95% CI, and percentages are the level of support that Δ*CA* > 0.

**Figure 4.**
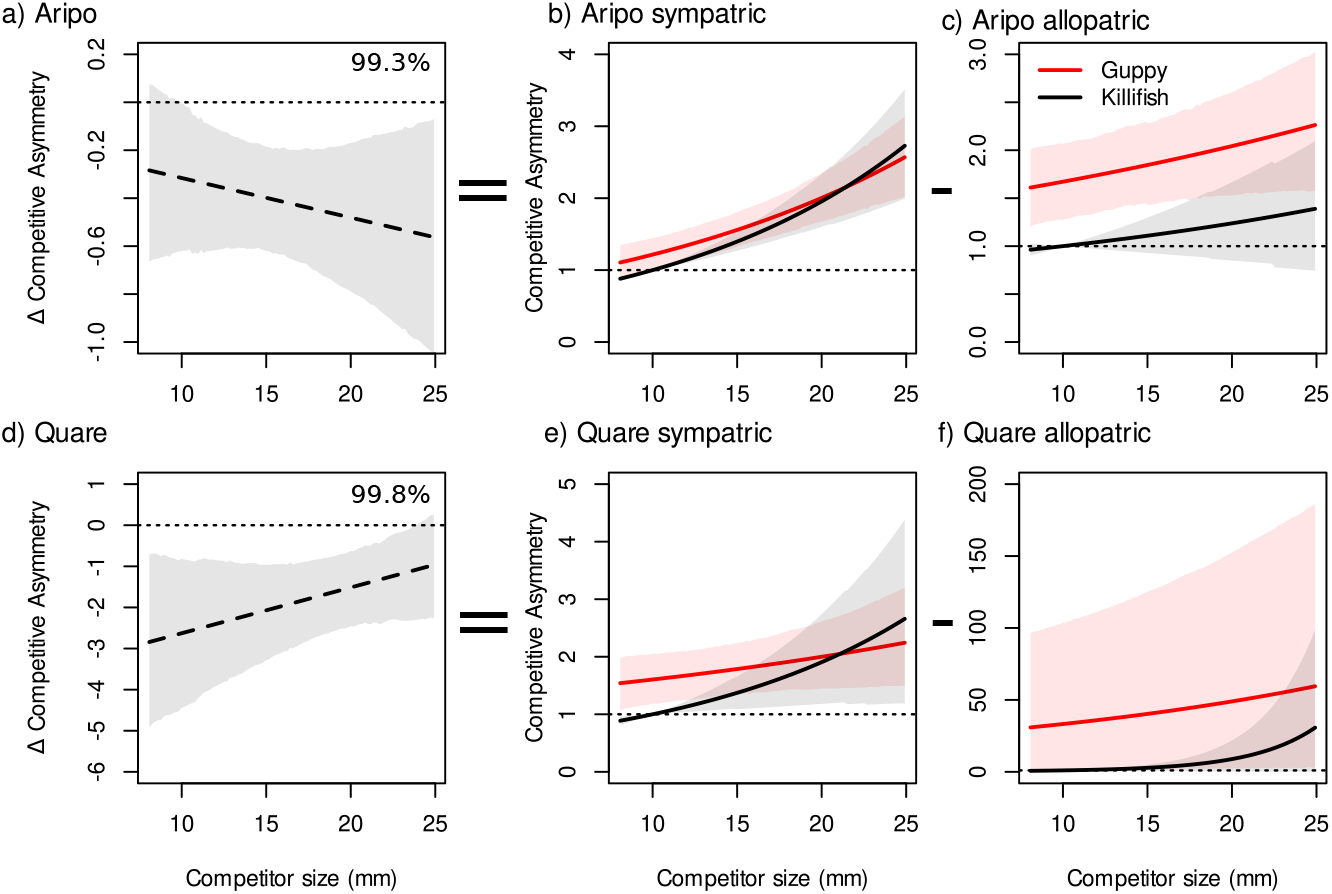
Change in competitive asymmetry for killifish. Panel a) shows the change in competitive asymmetry from allopatry to sympatry. Panels b), c), e), and f) show slices of the interaction surface for a “focal” 10mm killifish competing against either other killifish (black) or guppies (red) of all sizes. Values less than 1 mean that the competitive asymmetry favours the 10*mm* guppy, and values greater than one means favours the competitor. Shaded areas represent the 95% CI, and percentages are the level of support that Δ*CA* > 0.

In both competitive settings, the competitive asymmetries across all sizes favoured guppies. First, guppies exerted larger competitive effects on conspecifics compared to those exerted by killifish on guppies across all but the largest body sizes (Fig. 3). Second, the effect of interspecific competition of guppies on killifish was greater than the effect of intraspecific competition between killifish (Fig. 4).

Although the direction of change in species-dependent competitive asymmetries (following guppy invasion, i.e., from allopatric to sympatric communities) was consistent, the magnitude of this change differed between drainages (Fig. 2). In both drainages, co-evolved, sympatric killifish were stronger competitors to guppies than the allopatric killifish, but this effect was stronger in the Quare drainage compared to the Aripo (Quare Δ*η_GK_* = 2.628 [0.81,4.434]; Aripo Δ*η_GK_* = 0.316 [0.017, 0.637]; Fig. 2b).

There was a difference between the drainages in how the association between body size and competitive effect changed between allopatric and sympatric phenotypes (Δ*ϕ_G_* = *ϕ_G,sym_* – *ϕ_G,allo_* and Δ*ϕ_K_* = *ϕ_K,sym_* – *ϕ_K,allo_*). In the Aripo drainage, competitive effects increased with body size to a greater extent in sympatric fish than in allopatric fish (Δ*ϕ_K_* = 0.05 [0.009,0.085]; Δ*ϕ_G_* = 0.03, [0.004,0.057]). By contrast, in the Quare drainage, there was no clear difference in how competitive effects changed with body size between sympatric and allopatric phenotypes (Δ*ϕ_K_* = −0.128, [−0.270, 0.072]; Δ*ϕ_G_* = −0.017, [−0.050, 0.072]).

### How much do these changes in competitive asymmetry shift predictions about coexistence in guppies and killifish?

The two species IPM model of Bassar, Travis & Coulson (2017b), when parameterised with the interaction surfaces from these experiments, suggests that coexistence is slightly more likely between the sympatric phenotypes of both species than the allopatric phenotypes (Fig. 5). In general, throughout most of the parameter space defined by the size-dependent niche shifts in each species, the model predicts that guppies will exclude killifish. However, the area in which coexistence is possible expands from the allopatric to the sympatric phenotype combinations (Fig. 5). In the Aripo and Quare drainages, the percentage of parameter space in which coexistence is predicted to increase from allopatry to sympatry (Aripo: 6.6% to 8.3%; Quare: 0% to 12.2%). In both drainages, the region where killifish can exclude guppies also expands (Aripo: < 1% to 19%; Quare: 0% to 2.5%). Coexistence between these species is predicted to occur above the diagonal, where killifish shift their resource use with increased body size to a greater degree than guppies.

**Figure 5.**
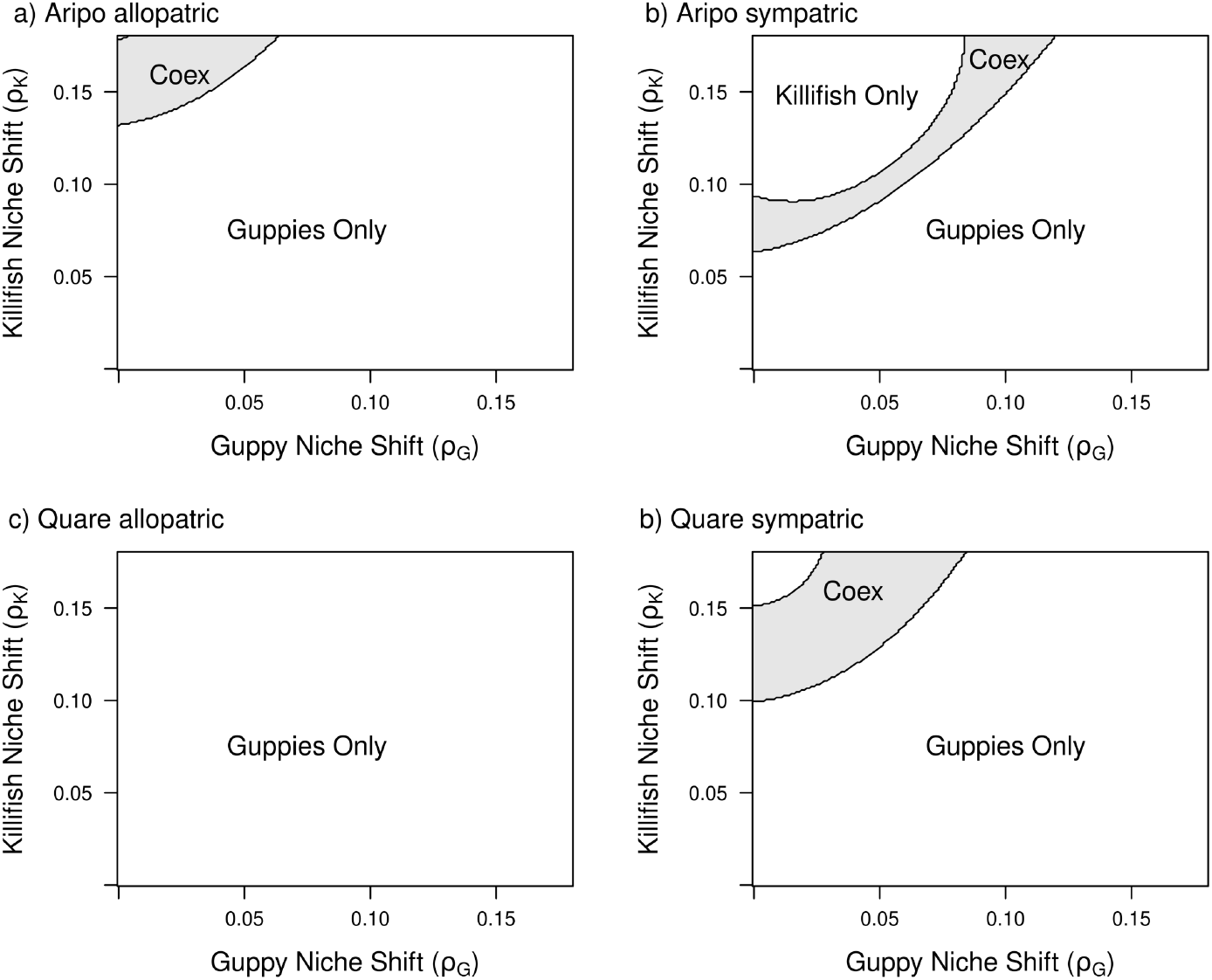
Predictions from IPM (Integral Projection Model) on species coexistence. The X- and y-axes represent how much guppies and killifish, respectively, shift their niches with increased body size. Light grey areas are those where guppies and killifish can coexist stably.

## Discussion

In this study, we quantified scenarios of intra and interspecific competition at two stages of the formation of a new fish community. We described the degree of competitive asymmetry between the initial (allopatric phenotypes) and final stages (sympatric phenotypes) of a guppy-killifish community formation, following the up-stream invasion of guppies into killifish habitats. In both stages of the KG community development, guppies tended to be stronger competitors than killifish. However, we found that guppies and killifish became more equal competitors in sympatry. This change in the degree of competitive asymmetry increased the likelihood of coexistence of these species in sympatry. The decrease in the competitive asymmetry between the allopatric and sympatric populations was due to changes in both species and size-dependent competitive asymmetries.

The different pattern of size-dependent competitive asymmetries between the two drainages suggests that there are several routes to achieve the same effect across the life cycle of guppies and killifish. In the Aripo drainage, the change in competitive asymmetry among small competitors was moderate compared to the change in larger competitors (Fig. 3A and 4A). In contrast, in the Quare drainage, the change in competitive asymmetry was much greater for smaller compared to larger competitors (Fig. 3D and 4D). Integrated across the entire life cycle of the guppies and the juvenile stages of killifish, the effect on the ability of each species to invade the other was similar; changes in competitive asymmetry led to an expanded region of coexistence and a region where killifish could exclude guppies, depending on the degree of ontogenetic niche shifts in each of the species (Fig. 5).

As predicted, the competitive advantage of guppies over killifish in allopatric and sympatric trials indicates that killifish should have stronger ontogenetic niche shifts than guppies in both newly founded and established KG communities (Balfour et al., 2003; Reichstein et al., 2013; Bassar *et al*., 2017a,b). There is some support for these predictions from niche occupancy from existing data. Guppies in KG communities have a broader resource use than guppies from KGP communities (Bassar *et al*., 2010; Zandona *et al*., 2011). Killifish rely more on terrestrial invertebrates than aquatic ones (Fraser *et al*., 1999; Owens, 2010) and it seems that as they grow larger, killifish increase their consumption of terrestrial prey (Dough Fraser *pers. comm.),* but currently there is no empirical data supporting this claim. Additionally, we do not currently know how differences in resource use at the species level or across sizes within a species map to the fitness of the two species.

In a previous study, Fraser & Lamphere (2013) found that allopatric (KO) killifish are stronger competitors than allopatric (KGP) guppies. In contrast to their study, we found that KGP guppies are better competitors than KO killifish. These differences are mainly the result of distinct experimental approaches. First, Fraser & Lamphere (2013) performed their experiment with similarly sized individuals of the two species, particularly guppies from the larger end of their natural size distribution. This restricts their results to only part of the range of sizes in which guppies and killifish partly share a niche, in particular the part in which competition for resources might not be so strong or where killifish may be stronger competitors than guppies. The range of body sizes used in this study captures most stages in the life history of guppies and the initial stages in the life history of killifish (Reznick *et al*., 2001; Walsh & Reznick, 2008), in which these species are more likely to compete for resources (Travis *et al.*, 2014). Our results are therefore a better representation of the effects of resource competition and allow us to derive more robust conclusions about the changes in competition among guppies and killifish in the region where their niches overlap.

A caveat to our study is that we performed the experiments with wild-caught individuals. Thus, a combination of phenotypic plasticity, experience, and genetic differences may each explain a significant part of the variation in competitive asymmetries of the fish used in this study. The ideal experiment should use second generation (F2) laboratory-bred individuals to separate the plastic and genetic differences underlying the differences in competitive ability and test the effects of competition on multiple life-history traits (e.g., survival and reproduction).

The evolution of size-competitive asymmetries increases the likelihood of coexistence, but only in relatively small areas of parameter space defining size-dependent niche shifts (Fig. 5). This suggests that KG guppies should still exclude KG killifish. Their demonstrable co-existence means that other ecological mechanisms must be acting. One possibility is that temporal or spatial variation may interact with size-structured interactions (e.g., competition and predation) to favour coexistence via storage effects and nonlinear effects competition on fitness (Warner & Chesson, 1985; Chesson, 2000; Kuang & Chesson, 2010). In the wild, guppies and killifish experience pronounced wet and dry seasons that change their resource landscape completely (Travis et al., 2014). During the wet season, aquatic prey items are drastically reduced while terrestrial prey items increase (Owens, 2010), a pattern that might offer a benefit to killifish. Indeed, guppy mortality is much higher in the wet season than the dry season, whereas the reverse is true for killifish (Travis *et al.*, 2014).

A second possibility is based on the fact that guppies and killifish each consume neonates of their own and the other species (Fraser & Gilliam, 1992; Fraser & Lamphere, 2013). The extent to which intraguild predation enhances the likelihood of coexistence depends on whether predatory effects are themselves symmetric between species (Bassar *et al.*, *unpublished data*). In this case, we do not yet know whether one species is a more voracious predator of the other and whether this component of the interaction evolves after initial contact as well.

Regardless of the mechanisms, this study demonstrates that the effects of body size on competition and niche differences between the species are likely to play an influential role in the evolution of coexistence. Size-based variation in these coexistence mechanisms is one way that individual differences can alter species coexistence through non-linear effects of competition on fitness (Hart *et al*., 2016). Given the ubiquity in nature of size-based interactions and their important ecological effects, understanding such interactions ought to be a priority for understanding coexistence (or the lack thereof) in natural communities.

## Acknowledgements

We would like to thank Jogi Ramlal and his family for providing housing and logistical support in Trinidad. We also like to thank the interns and graduate students at the Guppy project for their help in the field and in the laboratory, particularly to Joshua Goldberg, Dalton Oliver, Keegan Rankin, and Dara Yiu. Funding was provided by an NSF grant (1556884) to JT, RDB, and DNR and faculty start-up funds to RDB.

## Author Contributions

RDB developed the models, and RDB and JT conceived of the experiments with input from DNR. JMA, TP, AB, SC, NF performed the experiments. JMA and RDB analysed the results. JMA, RDB, and JT wrote the manuscript, and all authors contributed to the final draft.

## Data Availability Statement

The full data and the code will be deposit in Dryad.

