## Supporting Information for "The evolution of size-dependent competitive interactions promotes species coexistence"

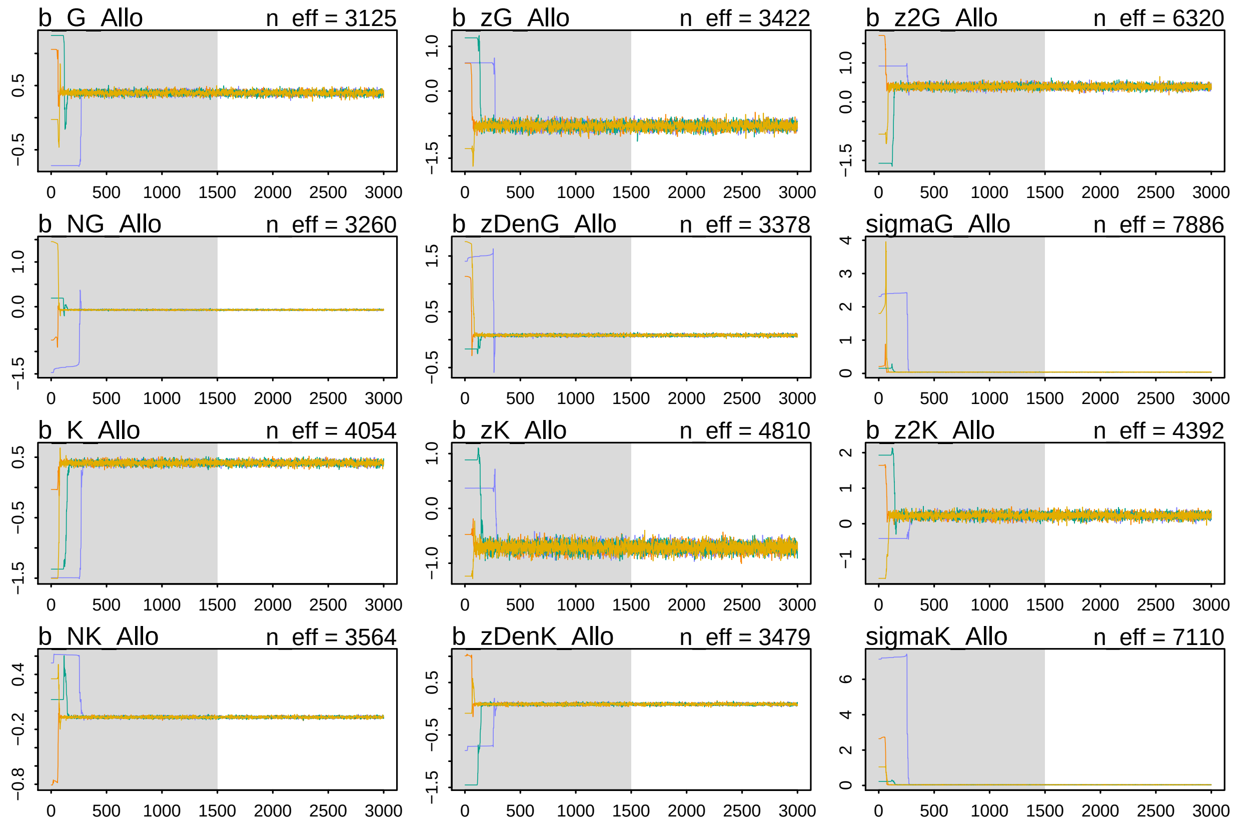

Figure-S1. Trace plots for the allopatric Aripo model showing that the four Hamilton Monte Carlo (HMC) chains converged for regression parameters .

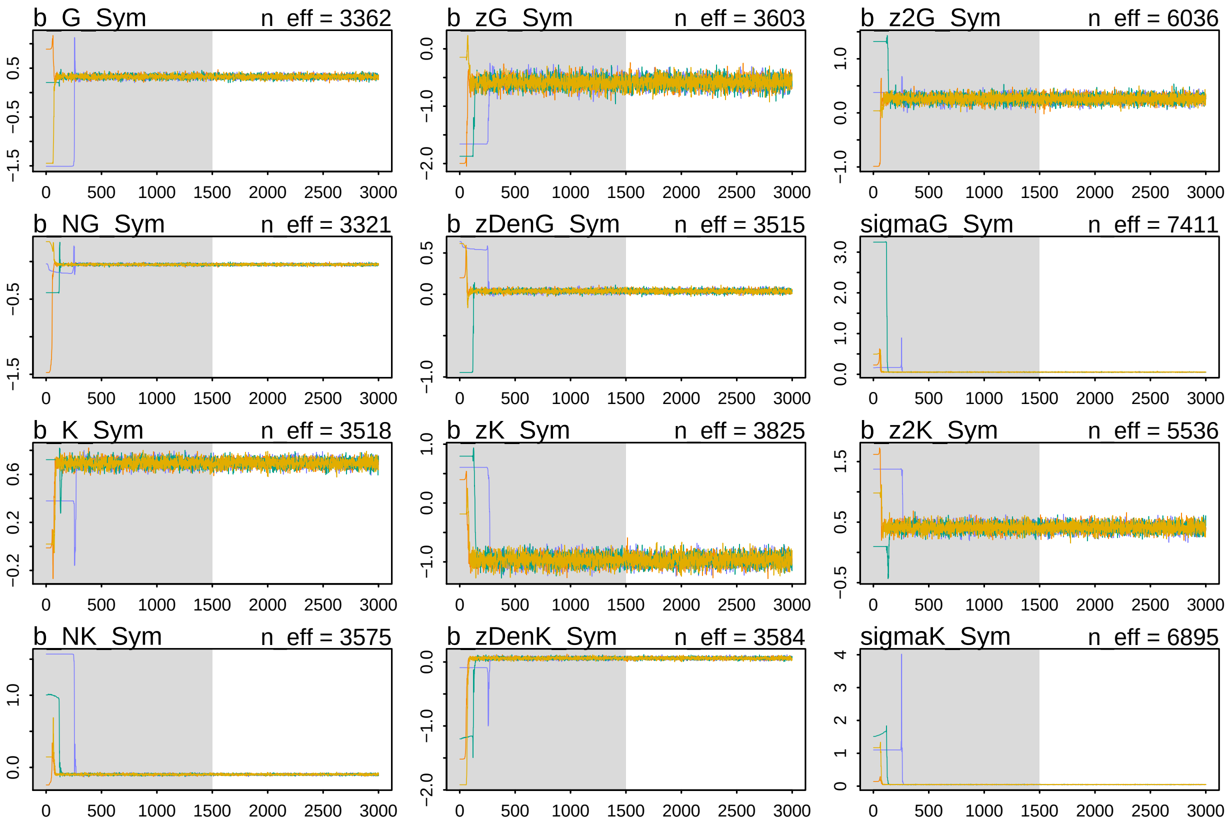

Figure-S2. Trace plots for the sympatric Aripo model showing that the four Hamilton Monte Carlo (HMC) chains converged for regression parameters.

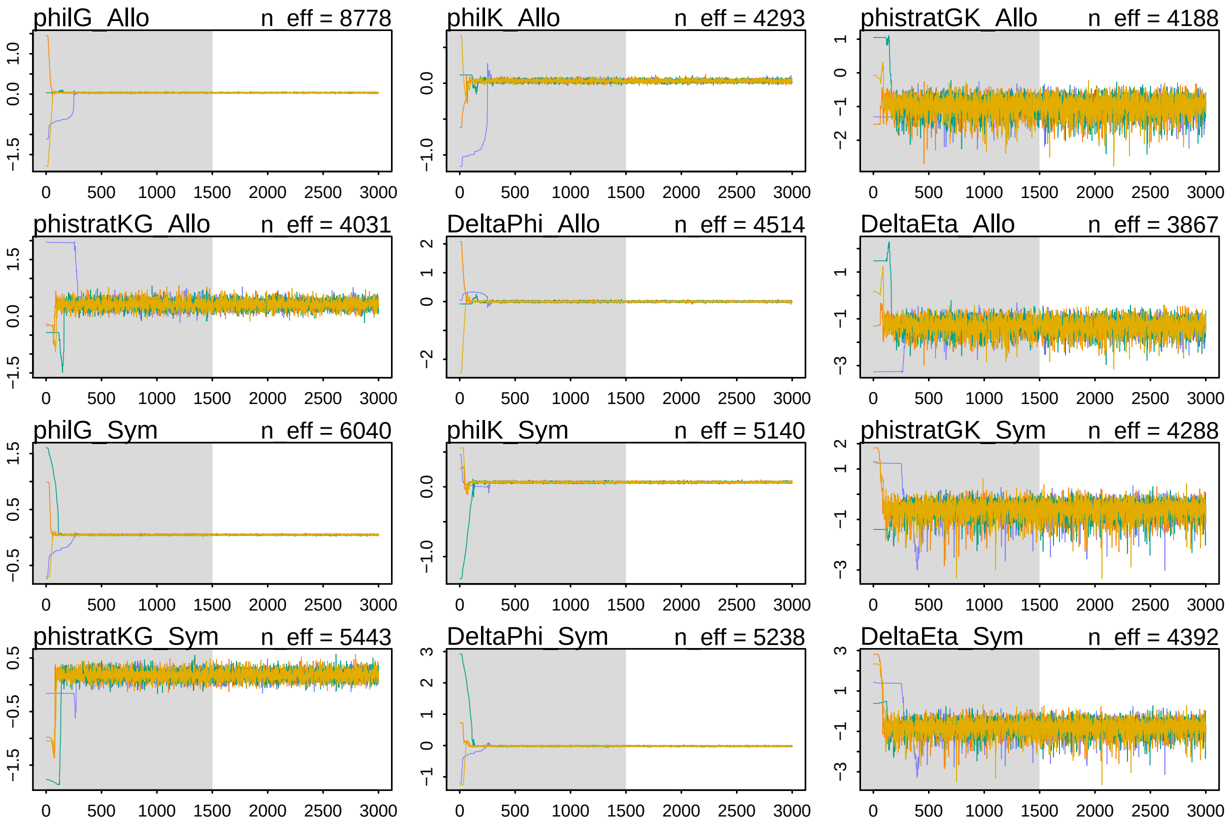

Figure-S3. TTrace plots for the Aripo model showing that the four Hamilton Monte Carlo (HMC) chains converged for interaction surface parameters.

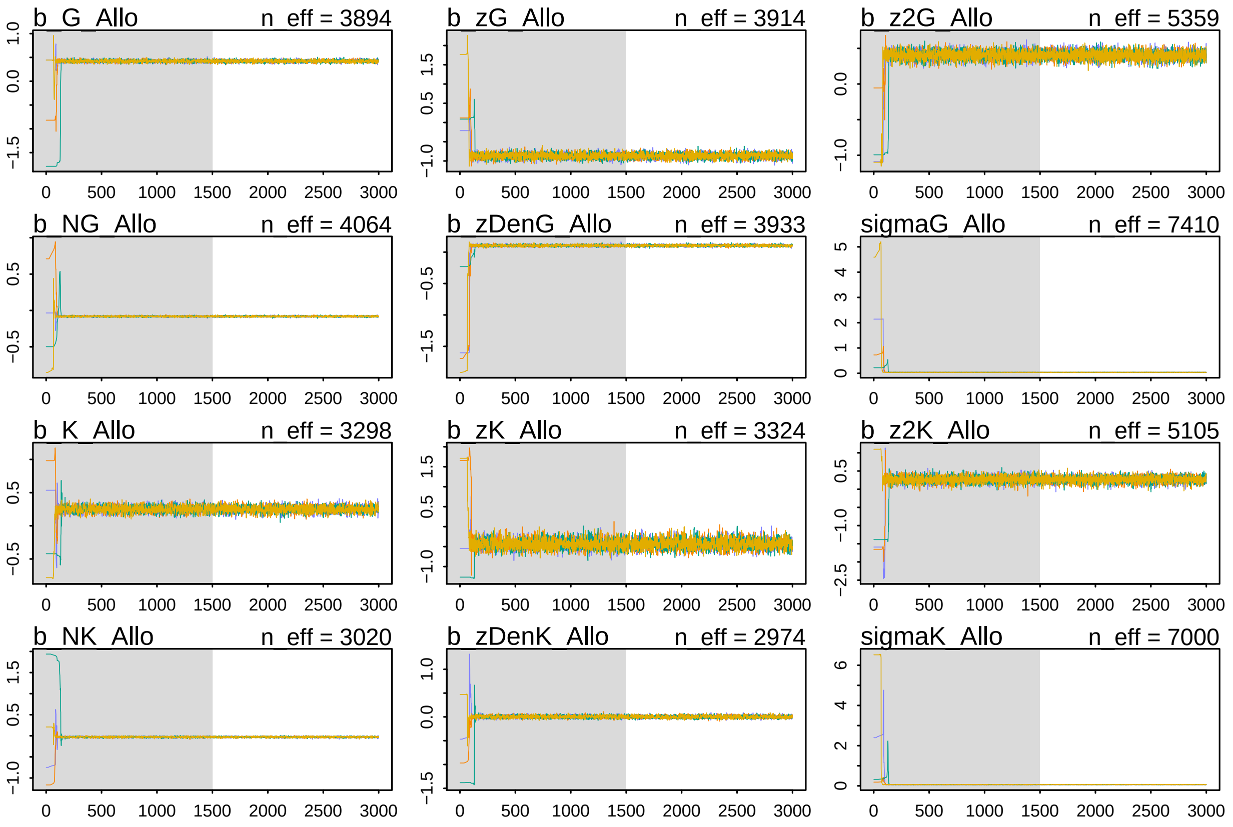

Figure-S4. Trace plots for the allopatric Quare model showing that the four Hamilton Monte Carlo (HMC) chains converged for regression parameters

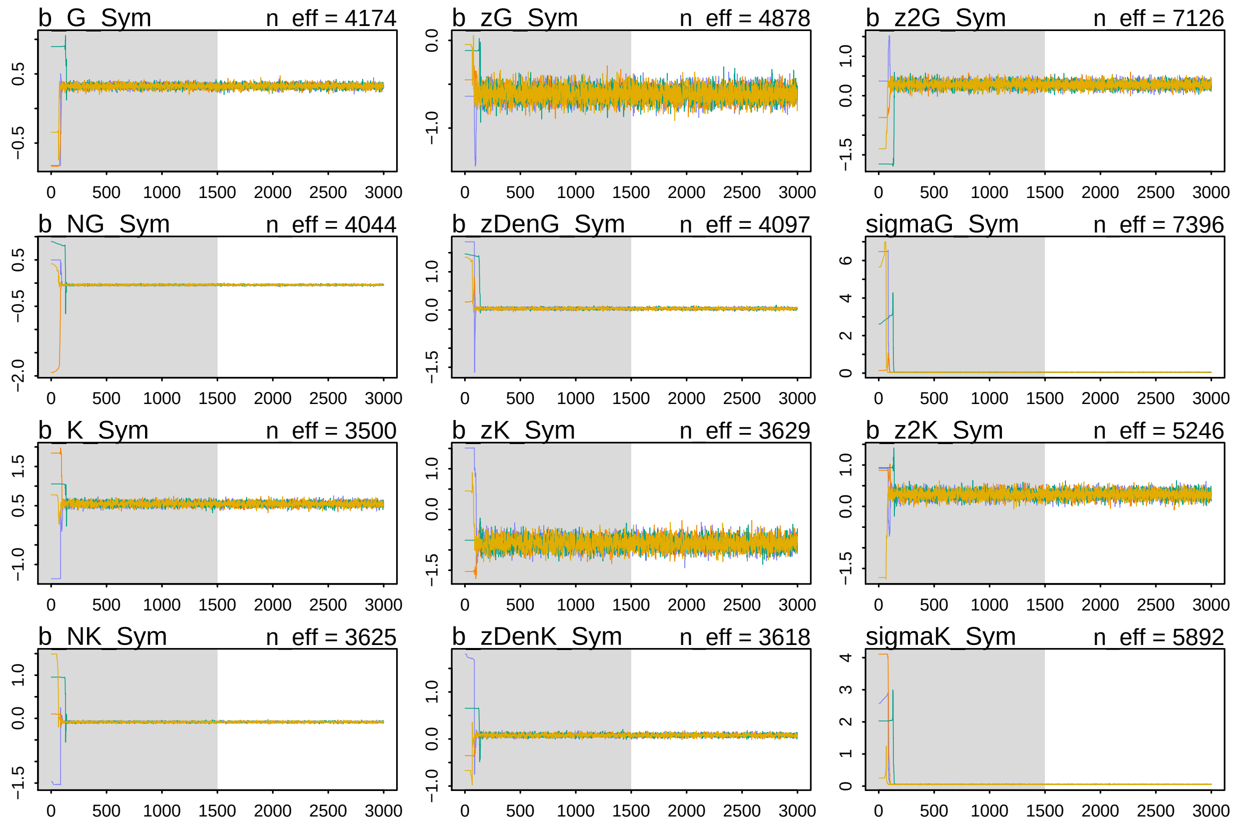

Figure-S5. Trace plots for the sympatric Quare model showing that the four Hamilton Monte Carlo (HMC) chains converged for regression parameters

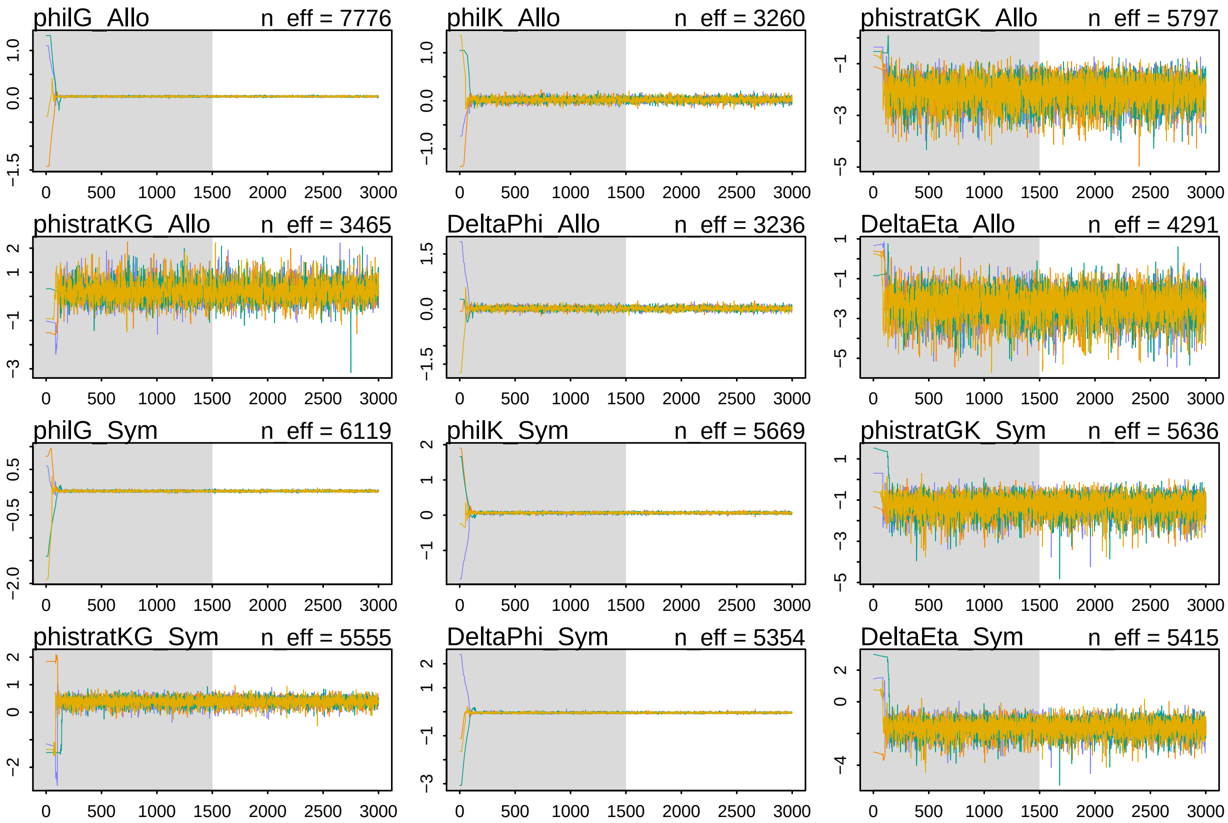

Figure-S6. Trace plots for the Quare model showing that the four Hamilton Monte Carlo (HMC) chains converged for interaction surface parameters

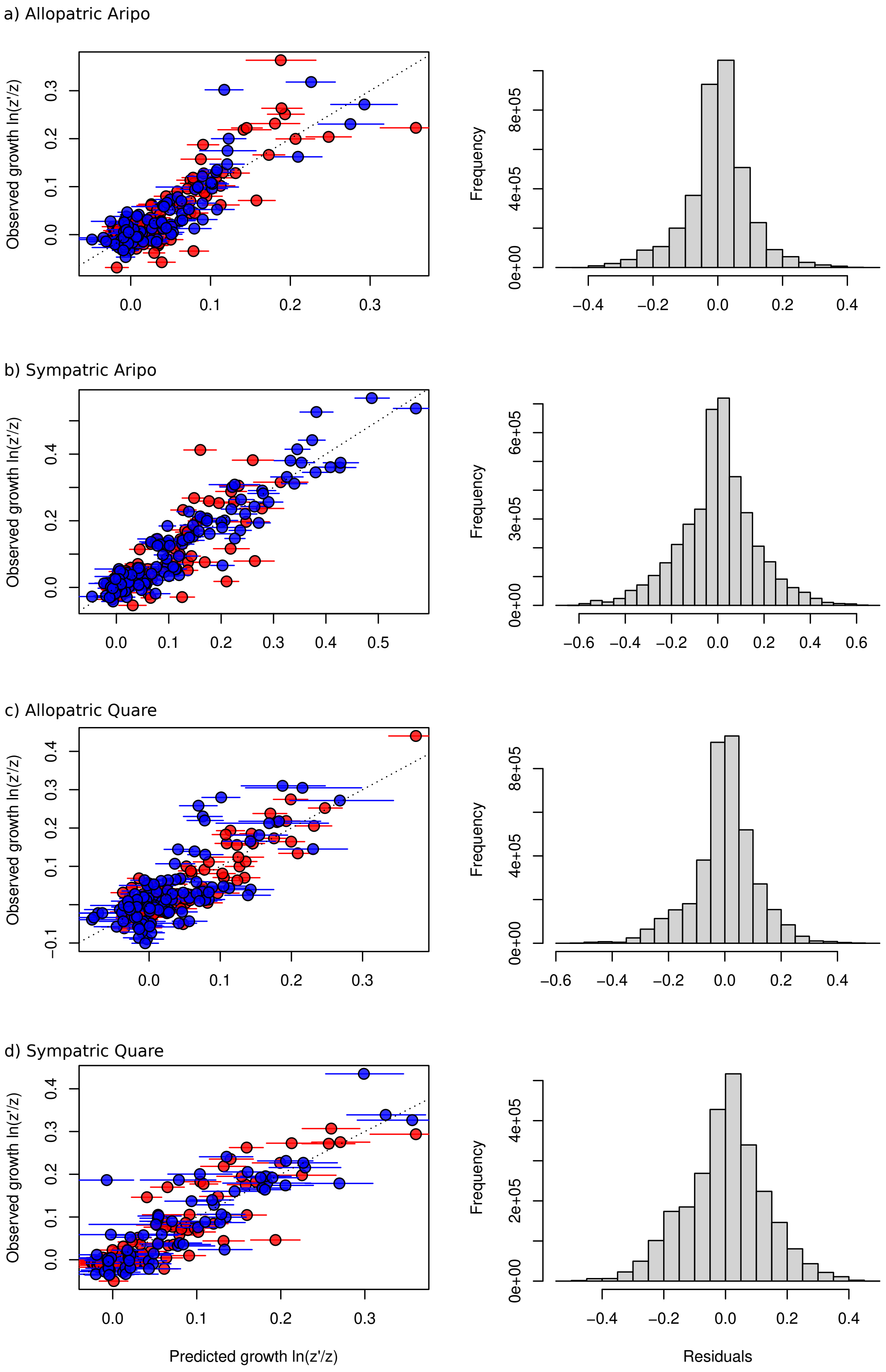

Figure-S7. Predicted vs Observed values and distribution of the model residuals. Circles and bars represent posterior predicted means (X-axis) and observed values (y-axis). Blue and red symbols represent guppies and killifish respectively.
